# Adaptation of turnip mosaic virus to *Arabidopsis thaliana* involves rewiring of VPg - host proteome interactions

**DOI:** 10.1101/2024.02.12.579887

**Authors:** José L. Carrasco, Silvia Ambrós, Pablo A. Gutiérrez, Santiago F. Elena

## Abstract

The outcome of a viral infection depends on a complex interplay between the host physiology and the virus, mediated through numerous protein-protein interactions. In a previous study we used high-throughput yeast two-hybrid (HT-Y2H) to identify proteins in *Arabidopsis thaliana* that bind to the proteins encoded by the turnip mosaic virus (TuMV) genome. Furthermore, after experimental evolution of TuMV lineages in plants with mutations in defense-related or proviral genes, most mutations observed in the evolved viruses affected the VPg cistron. Among these mutations, D113G was a convergent mutation selected in many lineages across different plant genotypes. In contrast, mutation R118H specifically emerged in the *jin1* mutant with affected jasmonate signaling. Using the HT-Y2H system, we analyzed the impact of these two mutations on VPg’s interaction with plant proteins. Interestingly, both mutations severely compromised the interaction of VPg with the translation initiation factor eIF(iso)4E, a crucial interactor for potyvirus infection. Moreover, mutation D113G, but not R118H, adversely affected the interaction with RHD1, a zinc-finger homeodomain transcription factor involved in regulating DNA demethylation. Our results suggest that RHD1 enhances plant tolerance to TuMV infection.

## 1. Introduction

Viruses establish multiple contact points with the host proteome, primarily through a limited number of proteins, particularly reduced in the case of RNA viruses (Belshaw, Pybus and Rambaut 2007; Mahmoudabadi et al. 2018). This interaction involves virus-host protein-protein interactions (PPIs) that result in the manipulation of various cellular pathways for creating favorable replication conditions. This manipulation can occur by sequestering cell resources for the virus’s benefit or by interfering with immune responses. Simultaneously, host cell receptors play a role in sensing foreign elements and responding accordingly. Recent years have seen significant progress in creating detailed, high-quality maps of virus-host PPIs (Uetz et al. 2006; de Chassey et al. 2008). Integrative approaches have identified both general and specific molecular mechanisms employed by different viruses (Dyer et al. 2008; Mukhtar et al. 2011; Pichlmair et al. 2012; Ahmed et al. 2018; Aguirre and Guantes 2023). These findings, coupled with additional omics data, have been used to predict phenotypic outcomes of infection (Tisoncik-Go et al. 2016; Cervera et al. 2018; Tarazona, Forment and Elena 2020). These advancements build upon the continuous development of a comprehensive large-scale map of host PPIs, which, while not fully complete, proves valuable in recognizing disease-associated modules (Mukhtar et al. 2011; Menche et al. 2015).

In a previous study Martínez et al. (2023) systematically identified direct PPIs between turnip mosaic virus strain YC5 (TuMV; species *Turnip mosaic virus*, genus *Potyvirus*, family *Potyviridae*) and one of its natural hosts, *Arabidopsis thaliana* (L.) HEYNH, using high-throughput yeast two-hybrid (HT-Y2H) screenings. This study uncovered 378 new PPIs between TuMV and plant proteomes. Among the viral proteins, the RNA-dependent RNA polymerase NIb emerged with the highest number of contacts, including crucial salicylic acid (SA)-dependent transcription regulators. The authors constructed and analyzed a network consisting of 399 TuMV-*A. thaliana* interactions, incorporating intravirus and intrahost connections. Notably, their findings revealed that TuMV-targeted host proteins (*i*) were enriched in various aspects of plant responses to infections, (*ii*) exhibit greater connectivity, (*iii*) had enhanced capacity to disseminate information throughout the cell proteome, (*iv*) showed higher expression levels, and (*v*) were under stronger purifying selection than expected by chance.

In previous evolution experiments with potyviruses, the VPg cistron has been identified as a significant target of selection. For instance, Agudelo-Romero *et al*. (2008) observed that a single amino acid replacement in VPg significantly increased tobacco etch virus (TEV) infectivity, severity of symptoms, and viral load in *A. thaliana*. Later on, Hillung et al. (2014) further described additional VPg mutants in several *A. thaliana* ecotypes that differed in their susceptibility to TEV infection. Likewise, Navarro et al. (2022) and Ambrós et al. (2024) evolved TuMV lineages in different defense-deficient genotypes of *A. thaliana*, observing pervasive mutations affecting VPg, with remarkable examples of parallel evolution with the same mutations fixing in lineages evolved in the same host genotype. Gallois et al. (2010) found that *A. thaliana* plants with knock-out mutations in the *EUKARYOTIC TRANSLATION INITIATION FACTORS (ISO)4E*, *(ISO)4G1* and *(ISO)4G2* (*eIF(iso)4E*, *eIF(iso)4G1* and *eIF(iso)4G2*, respectively) genes were resistant to TuMV infection, demonstrating the proviral role of these eIFs. However, two mutations in VPg were sufficient to overcome this resistance and revert to the original infection phenotype. Notably, Y2H assays showed that these mutations did not preclude the binding of VPg to eIF(iso)4E (Gallois et al. 2010). As another example, the *pvr2* locus, one of the most extensively used resistance genes against potato virus Y (PVY) in pepper cultivars, encodes the eIF4E factor, which physically interacts with VPg. Interestingly, all resistance-breaking viral isolates identified so far contain mutations in the VPg cistron (Duprat et al. 2002; Moury et al. 2004; Ayme et al. 2006).

VPg has been described as a scaffolding protein that interacts with other potyviral proteins (Bosque et al. 2014; Hafrén, Löhmus and Mäkinen 2015) and with many host proteins (Martínez et al. 2016, 2023), most notably factors involved in genome transcription, being linked to the 5’-end of the viral genome and providing the hydroxyl group that primes the synthesis of complementary strands by the viral NIb replicase (Eskelin et al. 2011). Additionally, it plays a role in protein synthesis by directly interacting with canonical translation factors eIF(iso)4E and eIF(iso)4G (Gallois et al. 2010; Hafrén, Löhmus and Mäkinen 2015) and with the antiviral factor eIF4E1 that protects the translational machinery during TuMV infection (Zafirov et al. 2023). Indeed, Martínez et al. (2023) HT-Y2H analyses and literature curation resulted in the identification of 43 direct interactors, involved in diverse cellular processes. Amongst the most highly expressed and connected genes, *CALNEXIN 1* (*CNX1*) and *OBERON 1* (*OBE1*) were shown to directly interact with VPg. Both CNX1 and OBE1 proteins were proviral factors whose knocked-out expression resulted in late and less severe symptoms. Other genes such as *NONEXPRESSER OF PATHOGENESIS-RELATED GENES 1* (*NPR1*), *SUMO 3* (*SUM3*), *SUMO CONJUGATING ENZYME 1* (*SCE1*), and specially *TGACG SEQUENCE-SPECIFIC BINDING FACTOR 1* (*TGA1*) also resulted in milder disease symptoms, although their interaction with VPg was mediated by NIb.

In various evolution experiments of TuMV isolate YC5 in different *A. thaliana* genotypes, it has consistently been observed that VPg fixed nonsynonymous mutations; 14 identified as being subject to positive selection (González, Butković, and Elena 2019; González et al. 2021; Navarro et al. 2022; Melero, González, and Elena 2023; Ambrós et al. 2024). The majority of these positively selected mutations are concentrated within a helix-coil-helix structural domain, spanning amino acids 110 to 120 [Fig. 4B in Ambrós et al. (2024)]. Certain mutations were consistently present across lineages that evolved in different host genotypes and environmental conditions. For instance, the VPg^D113G^ mutation was observed in lineages evolved in nine distinct plant genotypes, including wild-type Col-0 plants, and two environmental conditions (González, Butković, and Elena 2019; González et al. 2021; Navarro et al. 2022; Ambrós et al. 2024). Conversely, some mutations were specific to particular host genotypes, such as the VPg^R118H^ mutation, found exclusively in lineages evolved in mutants of *JASMONATE INSENSITIVE 1* (*JIN1*) and *DECREASED DNA METHYLATION 1* (*DDM1*) genes, both being particularly permissive to TuMV infection (Navarro et al. 2022; Ambrós et al. 2024).

**Figure 4.**
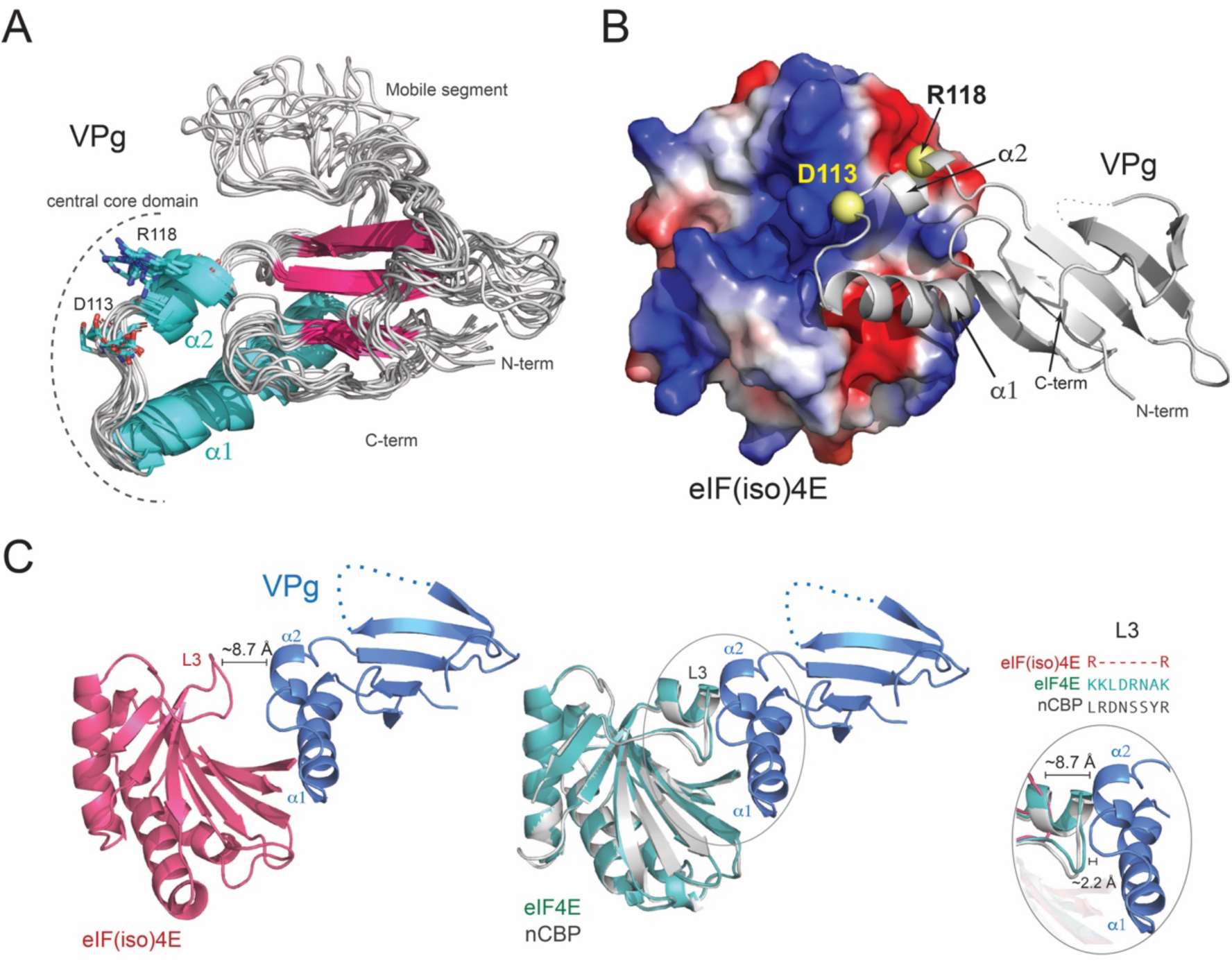
Structural analysis of the *A. thaliana* eIF(iso)4E - TuMV VPg complex. (A) The model of TuMV VPg reveals that D113 and R118 project outwards in a segment comprising the α1-α2 loop and helix α2. The structural ensemble illustrates the dynamical properties of the VPg proteins as determined by molecular dynamics simulations. (B) Analysis of the eIF(iso)4E - VPg docking model suggests that residues D113 and R118 bind to pockets of complementary charges in eIF(iso)4E. The reduced affinity of the D113G and R118H variants probably results from the removal of these ionic interactions. (C) The eIF(iso)4E - VPg (left) docking model explains the resistance in *eif(iso)4e* plants as the paralogous protein eIF4E and nCBP have a bulkier VPg binding region (L3) that is likely to impede binding to TuMV VPg (center). The sequence of the L3 region of the eIF4E homologs in *A. thaliana* and their relative distance to VPg is illustrated in the right panel.

Given the prominent role played by VPg during potyvirus infections, its apparent prominence as target of natural selection and its promiscuous interaction with other potyviral and host proteins, a question that immediately arises from all previously mentioned studies is the extent to which these mutations affect the ability of VPg to engage into PPIs resulting in fitness increases. To tackle this question, we have selected mutants VPg^D113G^ and VPg^R118H^ as examples of what can be considered generalist and specialist versions of VPg, respectively. The two VPg mutants were cloned and used as bait proteins in HT-Y2H experiments to identify host interactors that differ from those obtained for VPg^wt^ by Martínez et al. (2023). Then, the relative affinity of the differential interactors was evaluated by a quantitative assay and compared with the VPg^wt^. To further characterize the effect of these mutations in TuMV performance, we inoculated *eif(iso)4e* and *ros1-associated homeodomain protein 1* (*rhd1*) mutant plants and characterized disease progression and viral accumulation. Finally, we performed *in silico* evaluations of the effect of the two mutations of interest in VPg folding and used docking and molecular dynamics techniques to evaluate their effect on the interaction with eIF(iso)4E and RHD1.

## 2. Materials and Methods

### 2.1. Plasmid construction

The mutated variants of the VPg cistron were amplified by PCR using Phusion High-Fidelity DNA polymerase (Thermo Scientific, Waltham MA, USA) according to manufacturer’s recommendations and using primers specifically designed for the In-Fusion cloning system (Takara Bio, Mountain View CA, USA). The amplified PCR fragments were recombined into the yeast bait two-hybrid vector pGBKT7 (Takara Bio, Kusatsu, Japan), previously digested with *Eco*RI and *Bam*HI, using the In-Fusion enzyme. Thus, translational fusions of all three VPg protein variants (bait protein) with the GAL4 DNA-binding domain were generated.

### 2.2. High-throughput yeast two-hybrid screening

To search for differential interactors among the original VPg protein and the two selected mutated variants, an *A. thaliana* Col-0 cDNA library (Takara Bio) transformed into the *Saccharomyces cerevisiae* Y187 strain (prey strain) was screened using the Matchmaker^®^ Gold Yeast Two-Hybrid System (Takara Bio). For that purpose, the three bait VPg proteins cloned in pGBKT7 were transformed in the yeast strain Y2HGold. Then, the Y187 prey strain was mated to each of the Y2HGold haploid bait strains expressing the wild-type and the mutated variants of VPg protein and plated on a double dropout medium (SD/-Leu/-Trp) containing 40 μg/mL X-α-Gal and 200 ng/mL aureobasidin A. Co-transformants displaying α-galactosidase activity were subjected to a second round of selection on a quadruple dropout medium (SD/-Leu/-Trp/-Ade/-His) also containing X-α-Gal. Plasmids pGBKT7-T-antigen, pGADT7-laminin C, and pGADT7-murine p53 (Takara Bio) were used as negative controls. cDNA inserts from positive yeast clones were amplified by colony PCR, digested with *Alu*I and analyzed by agarose gel electrophoresis to eliminate duplicate clones. Selected prey plasmids were rescued using a glass-bead extraction protocol and transformed into *Escherichia coli* DH5α for amplification and sequencing. DNA and protein sequence analyses were performed with the WU-BLAST algorithm. For each positive interaction, the prey plasmid was re-transformed into the yeast strain Y2HGold and mated to all three bait strains and also to a control Y187 yeast strain transformed with the empty pGBKT7 vector to remove false positives.

### 2.3. Plant genotypes, virus clones and inoculation procedures

Wild-type and two *A. thaliana* mutants were used in this study, all in the same Col-0 background. Seeds of the T-DNA insertion line SALK_092897C (*rhd1-1* mutant) were obtained from the Nottingham Arabidopsis Stock Center (NASC), whereas *eif(iso)4e* seeds were a kind gift from Dr. Jean-Luc Gallois (INRA, Avignon, France) (Duprat et al. 2002).

TuMV infectious sap was obtained from TuMV-infected *Nicotiana benthamiana* DOMIN plants inoculated with the infectious plasmid p35STunos that contains a cDNA of the TuMV genome isolate YC5 from calla lily (*Zantesdeschia* sp) (GeneBank AF530055.2) under the control of the cauliflower mosaic virus 35S promoter and the *nos* terminator (Chen et al. 2003) as described elsewhere (González, Butković and Elena 2019; Corrêa et al. 2020). After plants showed symptoms of infection, they were pooled and frozen with liquid N_2_. This frozen plant tissue was homogenized into a fine powder using a Mixer Mill MM400 (Retsch GmbH, Haan, Germany). For inoculations, 0.1 g of powder was diluted in 1 mL inoculation buffer [50 mM phosphate buffer (pH 7.0), 3% PEG6000, 10% Carborundum] and 5 μL of the inoculum was gently rubbed onto two leaves per plant. As a control, another set of plants were mock-inoculated only with inoculation buffer. Plants were all inoculated when they reached growth stage 3.5 in the Boyes et al. (2001) scale. This synchronization ensures that all hosts were at the same phenological stage when inoculated.

VPg mutations D113G and R118H were introduced by site-directed mutagenesis in the p35STunos plasmid using the Quikchange II XL kit (Agilent Technologies, Santa Clara CA, USA) and following the indications of the manufacturer, resulting in plasmids p35STunos-VPg^D113G^ and p35STunos-VPg^R118H^. Plasmids were purified using the ZymoPURE II Plasmid Miniprep kit (Zymo Research, Irving CA, USA) and resuspended in sterile water.

For evaluation of the mutated VPg alleles, wild-type, *eif(iso)4e* and *rhd1-1* plants were inoculated with p35STunos, p35STunos-VPg^D113G^ and p35STunos-VPg^R118H^. One leaf per plant was inoculated by gentle abrasion after applying 3 μg of plasmid DNA diluted in 6 μL of water supplemented with 10% Carborundum.

### 2.4. Infectivity, symptoms progression and quantification of viral load

The number of infected symptomatic plants, out of 20, as well as the severity of symptoms in a semi-quantitative scale in which zero means no symptoms and 5 systemic necrosis [Fig. 1 in Butković et al. (2021)] were monitored every day until 14 days post-inoculation (dpi). As a control, another set of mock-inoculated plants was treated in the same way.

**Figure 1.**
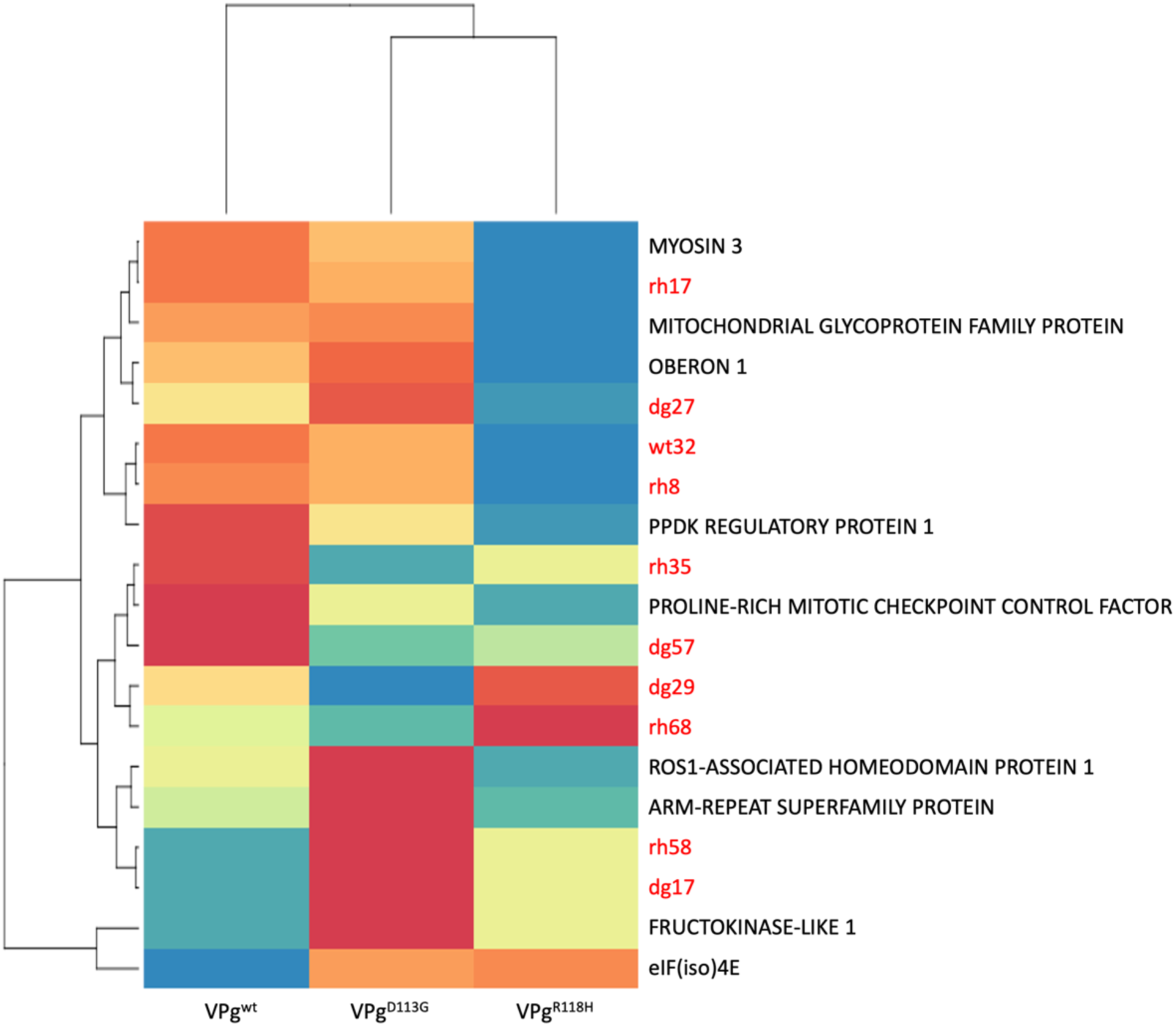
Heatmap constructed from the estimated binding affinities between the three different VPg alleles (wild-type, D113G and R118H) and the 19 host proteins identified in the HT-Y2H screening. Gene names are only provided for interactors for which statistically significant differences with VPg^wt^ were found and further identified by sequencing. Interactors for which no significant differences in binding affinity were observed are indicated in red. Labels starting with “dg” correspond to those found in the screening against VPg^D113G^, “rh” to those found in the screening against VPg^R118H^ and “wt” against the VPg^wt^ allele. Color scale: the more intense blue corresponds to stronger affinities than for VPg^wt^, whereas stronger red corresponds to weaker affinities than for VPg^wt^.

Viral loads were measured 12 dpi in four different pools of five infected plants each, by absolute real-time quantitative RT-PCR (RT-qPCR) using a TuMV standard curve and the primers TuMV F117-fw (^5’^CAATACGTGCGAGAGAAGCACAC_3’_) and F118-rv (^5’^TAACCCCTTAACGCCAAGTAAG_3’_) that amplify a 173 nucleotides fragment from the CP cistron of TuMV genome, as previously described (Corrêa et al. 2020). Briefly, standard curve consisted of nine serial dilutions of the *in vitro* synthesized TuMV genome prepared in total plant RNA purified from healthy *A. thaliana* plants. Amplification reactions were run in a 20 μL volume using the qPCRBIO SyGreen 1-step Go Hi-ROX System (PCRBiosystems, London, UK) and the recommended manufacturer’s instructions in an ABI StepOne Plus Real-time PCR System (Applied Biosystems, Foster City CA, USA). An initial RT phase of 10 min at 45 °C and 2 min at 95 °C was followed by a PCR stage with the following cycling conditions: 40 cycles of 5 s at 95 °C and 25 s at 60 °C, and a final melting curve profile analysis that consisted of 15 s at 95 °C, 1 min at 60 °C and 15 s at 95 °C. As negative controls, total plant RNA (noninfected control) and water were included in the analysis. Quantitative reactions were run as three technical replicates per sample and results were analyzed using the StepOne software 2.2.2 (Applied Biosystems).

### 2.5. Determination of TuMV consensus sequence

The consensus sequences of the viral genomes accumulated at the end of the infection processes were generated as described elsewhere (*e.g*., Navarro et al. 2022). In short, total RNAs were obtained and amplified by high-fidelity RT-PCR using the AccuScript Hi-Fi (Agilent Technologies) reverse transcriptase and Phusion DNA polymerase (Thermo Scientific) following the manufacturer’s instructions. Each complete TuMV genome was amplified as three overlapping amplicons using three specific sets of primers and the amplification conditions described in Navarro et al. (2022). PCR products were purified with the MSB Spin PCRapace kit (Stratec Molecular, Coronado CA, USA) and then Sanger-sequenced by Novogene Europe Co. Ltd. (Cambridge, UK). Full-length consensus viral sequences were obtained assembling the sequences of the three amplified products by using the Genious R9.0.2 program (Dotmatics, Bishop’s Stortford, UK). No additional mutations were found in any of the VPg clones.

### 2.6. Host gene expression analysis by relative RT-qPCR

For the analysis of *RHD1* and *TGA1* expression, RNA was purified with the Quick-RNA Plant Miniprep kit (Zymo Research) following the protocol recommended by the manufacturer. RT-qPCR was performed in an ABI OneStep Plus Real-time PCR System (Applied Biosystems) from 50 ng RNA using the qPCRBIO SyGreen 1-step Go Hi-ROX System from (PCRBiosystems). The *AT1G13320* corresponding to the *PROTEIN PHOSPHATASE 2A SUBUNIT A3* (*PP2AA3*) gene was used as endogenous reference (Czechowski et al. 2005). The primers used for the amplification were pp2aa3-fw (^5’^TTGGTGCTCAGATGAGGGAGAG_3’_), pp2aa3-rv (^5’^TTCACCAGCTGAAAGTCGCTTAG_3’_), rhd1-fw (^5’^TATCGGAACCACCGCTTACG_3’_), rhd1-rv (^5’^CGGTGGAAGTTACGGTGACA_3’_), tga1-fw (^5’^ACGAACCTGTCCATCAATTCGG_3’_), and tga1-rv (^5’^CCATGGGAAGTATCCTCTGACACG_3’_). For expression in leaves, plants were grown for about 4 weeks [growth stage ∼3.70 in the Boyes et al. (2001) scale].

### 2.7. Statistical analyses

Quantitative specificity α-galactosidase assays were conducted per interactor and VPg allele, with repetitions ranging from 1 to 5 times (median 2) in independent blocks. To mitigate potential block effects, individual measures were normalized by the mean of the VPg^wt^ allele estimated in the corresponding block. Normalized data underwent analysis using Welch’s robust one-way ANOVA, and effect magnitudes were assessed using the *η*^2^ statistic. Conventionally, *η*^2^ < 0.05 are considered small, 0.05 ≤ *η*^2^ < 0.15 as medium, and *η*^2^ ≥ 0.15 as large effects.

Relative expression data for *RHD1* and *TGA1* were fitted to a generalized linear model (GLM) with infection status (mock-inoculated *vs*. TuMV-VPg^wt^ infected) and days post-infection (dpi) as orthogonal factors, utilizing a Gamma distribution and log-link function. In this context, the magnitude of effects was assessed using the 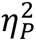, with similar criteria than for *η*^2^ above.

The number of infected plants, out of 20, observed for wild-type and *rhd1* plants were independently analyzed using Kaplan-Meier regression. Pairwise comparisons of the median time to the appearance of first symptoms for each VPg allele were performed using log-rank tests. Symptom severity progression data were independently fitted to repeated-measures ANOVAs, with dpi as an intra-individual factor and VPg allele as an inter-individual factor. The magnitude of effects was evaluated using the 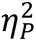 statistic.

Viral load data were fitted to a GLM with plant genotype and VPg allele as orthogonal factors, and experimental block nested within the interaction of the two orthogonal factors, employing a Gaussian distribution and identity-link function. The magnitude of effects was assessed using the 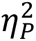 statistic.

In all reported pairwise *post hoc* tests, the Bonferroni sequential method was applied.

### 2.8. Homology modelling

Models were generated with MODELLER version 10.4 (Webb and Sali, 2016). For the *A. thaliana* eIF(iso)4E, and its paralogs eIF4E and nCBP, the X-ray structures of eIF4E from *Cucumis melo* (Miras et al. 2017), *Pisum sativum* (Ashby et al. 2011), and *Mus musculus* (Rydzik et al. 2017; Wan et al. 2020; Wojcik et al. 2021) were used as templates (Supplementary Fig. S1). The model of the VPg protein from TuMV was inferred using the NMR structure of the PVY homolog (Coutinho de Oliveira et al. 2019). Input alignments were generated with MUSCLE (Edgar, 2004). In each case, the model lowest-energy model out of 100 structures was selected and submitted to a final round of refinement using ROSETTA’s relax protocol (Conway et al. 2013).

### 2.9. Docking

Docking was performed with the Monte Carlo based multi-scale docking algorithm implemented in RosettaDock (Chaudhury et al. 2011). In each case, high-energy rotamers were removed prior to docking using the Docking Prepack application from ROSETTA (Wang et al. 2005). For the eIF(iso)4E and VPg docking model, structures were prepositioned near each other with the binding pockets facing each other using HADDOCK (Dominguez et al. 2003; van Zundert et al. 2016; Honorato et al. 2021). In this case, ambiguous interaction restraints (AIRs) were generated using a combination of NMR chemical shift perturbations (Coutinho de Oliveira et al. 2019), mutagenesis data on the eIF4E/eIF(iso)4E - VPg system (Monzingo et al. 2007; Roudet-Tavert et al. 2007; German-Retana et al. 2008; Ashby et al. 2011; Pérez et al. 2012; Svanella-Dumas et al. 2014), and experimental data from Martínez et al. (2023). Unfolded and mobile regions were removed from both proteins and docking site constraints for residues known to be in contact in the eIF(iso)4E/VPg interaction were included in local docking protocol (Supplementary Fig. S2). A total of 1000 decoys were generated during the docking perturbation runs. Residues in the interface of the selected model were refined using a high-resolution full atom minimization protocol (Conway et al. 2013). After this step, the mobile regions of VPg and eIF(iso)4E were included using MODELLER, and an optimal conformation was recalculated using the kinematic closure method in ROSETTA (Mandell et al. 2009) (Supplementary Fig S2). The final complex was further refined using a molecular dynamic simulation in explicit solvent with NAMD (Phillips et al. 2020). This step was performed using periodic boundary conditions in a water box with a 10 Å padding, 0.15 M NaCl, and a temperature of 298 K (Spivak et al. 2023). Docking of RHD1 and the VPg protein was performed using a global docking protocol using the AlphaFold RHD1 model available at Uniprot (Jumper et al. 2021). Docking was performed separately on N- and C-terminal Zn-finger and homeobox domains, respectively. After minimization, structures were fitted using a global docking protocol using 10,000 decoys for each RHD1 domain. Other RHD1 PPIs were pulled out from STRING database version 12.0 (Szklarczyk et al. 2023).

## 3. Results and Discussion

### 3.1. HT-Y2H screens and evaluation of differences in affinity between VPg variants and host interactors

Three screens were conducted in parallel using wild-type and two mutated VPg variants, D113G and R118H, as the respective bait proteins. The number of clones screened ranged from 2·10^7^ to 2.5·10^7^ per variant, with a mating efficiency of 30% to 35%. Initially, approximately 400 clones were selected, from which six for VPg^wt^, seven for VPg^D113G^, and eight for VPg^R118H^ were further validated through a second round of more rigorous selection and semiquantitative evaluation of binding specificity after plasmid rescue and yeast retransformation. Subsequently, a quantitative α-galactosidase activity assay was performed with the selected clones. Only clones displaying significant statistical differences among alleles were subjected to gene identification through sequencing. Following sequencing, two clones selected for VPg^D113G^ corresponded to *AT5G42780*, while two clones selected for VPg^wt^ matched with *AT5G35620*. Consequently, we identified 19 host proteins that interact differentially among the three VPg alleles.

Fig. 1 depicts a heatmap constructed based on the binding affinities similarity matrix among the 19 host proteins and the three VPg alleles. Among the interactors, nine exhibited a significant overall difference among the three VPg alleles (Supplementary Fig. S3). The smallest significant effect was observed with *AT2G19270*, which encodes the *PROLINE RICH MITOTIC CHECKPOINT CONTROL FACTOR* (*PRCC*). PRCC is involved in regulating protein modifications and signal transductions (Welch’s robust ANOVA: *F*_2,25.451_ = 4.326, *P* = 0.024, *η*^2^ = 0.171). On the other hand, the most substantial effect was observed with *AT5G35620*, the locus encoding the well-characterized VPg interactor eIF(iso)4E protein (*F*_2,67.604_ = 675.852, *P* < 0.001, *η*^2^ = 0.958).

Applying Bonferroni *post hoc* pairwise difference tests, we compared the affinity values between VPg^wt^ and each of the 19 host interactors with those observed for the two VPg mutant alleles, revealing a remarkably diverse pattern of effects. As illustrated in Fig. 1, these 19 proteins can be categorized into three distinct groups. In the following discussion, we will delve into the host proteins exhibiting differential binding strength with the VPg alleles. The first group comprises two proteins whose interactions are significantly more robust with VPg^wt^ than with either of the two mutant alleles. These proteins are: (*i*) the aforementioned eIF(iso)4E (*P* < 0.001; 90.50% weaker interaction) and (*ii*) the FRUCTOKINASE-LIKE 1 (FLN1) protein, a member of the pfkB-carbohydrate kinase family encoded by locus *AT3G54090* (*P* < 0.001; 36.54% weaker interaction). FLN1 serves as a potential plastidial target of thioredoxin z and is crucial for proper chloroplast development. Its role extends to the regulation of plastid-encoded polymerase (PEP)-dependent chloroplast transcription. Consequently, *fln1* mutants exhibit aberrant chloroplast development, general developmental defects, and impaired PEP-dependent transcription (Gilkerson et al. 2012).

The second group in Fig. 1 is the most heterogeneous, encompassing three host proteins that exhibit either no differences with VPg^wt^ or variable effects with VPg^D113G^ and VPg^R118H^. (*i*) The *AT5G62580* locus encodes ARM-REPEAT SUPERFAMILY PROTEIN and demonstrates a 28.76% reduction in binding strength with VPg^D113G^ compared to VPg^wt^ (*P* = 0.025), while it shows no significant effect on VPg^R118H^ (*P* = 1.000). This family of proteins binds to microtubules, impacting cellular organization and organ growth (Buschmann et al. 2004). Intriguingly, Martínez et al. (2023) described this protein as an interactor of NIb replicase. (*ii*) The second protein in this group is RHD1 (encoded by *AT5G42780*). RHD1, a zinc finger and homedomain (ZF-HD) protein, interacts with ROS1-ASSOCIATED METHYL-DNA BINDING PROTEIN 1 (RMB1), ROS1-ASSOCIATED WD40 DOMAIN-CONTAINING PROTEIN (RWD40) and REPRESSOR OF SILENCING 1 (ROS1) in a multiprotein complex involved in the base excision repair pathway through DNA methylation. This interactor was not described by Martínez et al. (2023). Interestingly, RHD1 interacts with the transcriptional regulator TGA1, known to act as a proviral factor via its interaction with NIb (Martinez et al. 2023). RHD1 exhibits a 22.62% decrease in binding affinity with VPg^D113G^ (*P* < 0.001) but a 17.35% increase with VPg^R118H^ (*P* = 0.003) compared to VPg^wt^. (*iii*) The third member of this group is the product of the *PRCC* gene, involved in the regulation of protein modifications and signal transductions. While VPg^D113G^ shows no differences in binding compared to VPg^wt^ (*P* = 1.000), VPg^R118H^ exhibits 22.5% stronger binding (*P* = 0.029).

The third group in Fig. 1 comprises four host proteins that exhibit no differences in affinity between VPg^wt^ and VPg^D113G^ but consistently show increased binding strength with VPg^R118H^. (*i*) The product of the gene *PYRUVATE ORTHOPHOSPHATE DIKINASE (PPDK) REGULATORY PROTEIN 1* (*RP1*) (*AT4G21210*) demonstrates 47.15% stronger binding with VPg^R118H^. RP1, a bifunctional serine/threonine kinase and phosphorylase, is involved in the dark/light-mediated regulation of PPDK by catalyzing its phosphorylation/dephosphorylation (Chastain et al. 2007). Notably, RP1 was not described by Martínez et al. (2023). (*ii*) The second host protein in this group is OBERON 1 (OBE1), encoded by locus *AT3G07780*; this interaction was previously described by Martínez et al. (2023). OBE1, a nuclear PHD finger protein, interacts with WRKY transcription factors, forming complexes that bind to histones (via the PHD finger) and repress the transcription of many stress-responsive genes (Du et al. 2023). (*iii*) The MITOCHONDRIAL GLYCOPROTEIN FAMILY PROTEIN encoded by locus *AT1G15870* binds 36.19% more strongly with VPg^R118H^ than with VPg^wt^. Unfortunately, no information about the function of this glycoprotein in the context of infection is currently available. (*iv*) The MYOSIN 3 (MYOS3) protein encoded by locus *AT3G58160*, a class XI myosin gene, is involved in the trafficking of Golgi stacks, peroxisomes, and mitochondria in root hairs and leaf epidermal cells (Peremyslov et al. 2008). Class XI myosin genes have been shown to play important and diverse roles in virus replication. For example, Amari et al. (2014) demonstrated that the inactivation of myosin XI genes in *Nicotiana benthamiana* affected the functioning of the endoplasmic reticulum, resulting in the aggregation of the movement protein of tobacco mosaic virus and the altered intracellular distribution of the viral replicase. MYOS3 binds to VPg^R118H^ 27.51% more strongly than to the VPg^wt^ allele (*P* = 0.007).

### 3.2. TuMV infection affects the expression of *RHD1* and *TGA1*

As outlined in the Introduction, VPg interacts with both the viral replicase NIb and the host factor RHD1. Additionally, both NIb and RHD1 interact with the transcriptional regulator TGA1, identified as a proviral factor by Martínez et al. (2023) for the first time. To assess the impact of TuMV infection on the expression of these crucial genes, a time-course analysis of *RHD1* and *TGA1* expression was conducted via relative RT-qPCR in three pools of mock- and TuMV-VPg^wt^-inoculated wild-type plants. Fig. 2 presents the results of these experiments.

**Figure 2.**
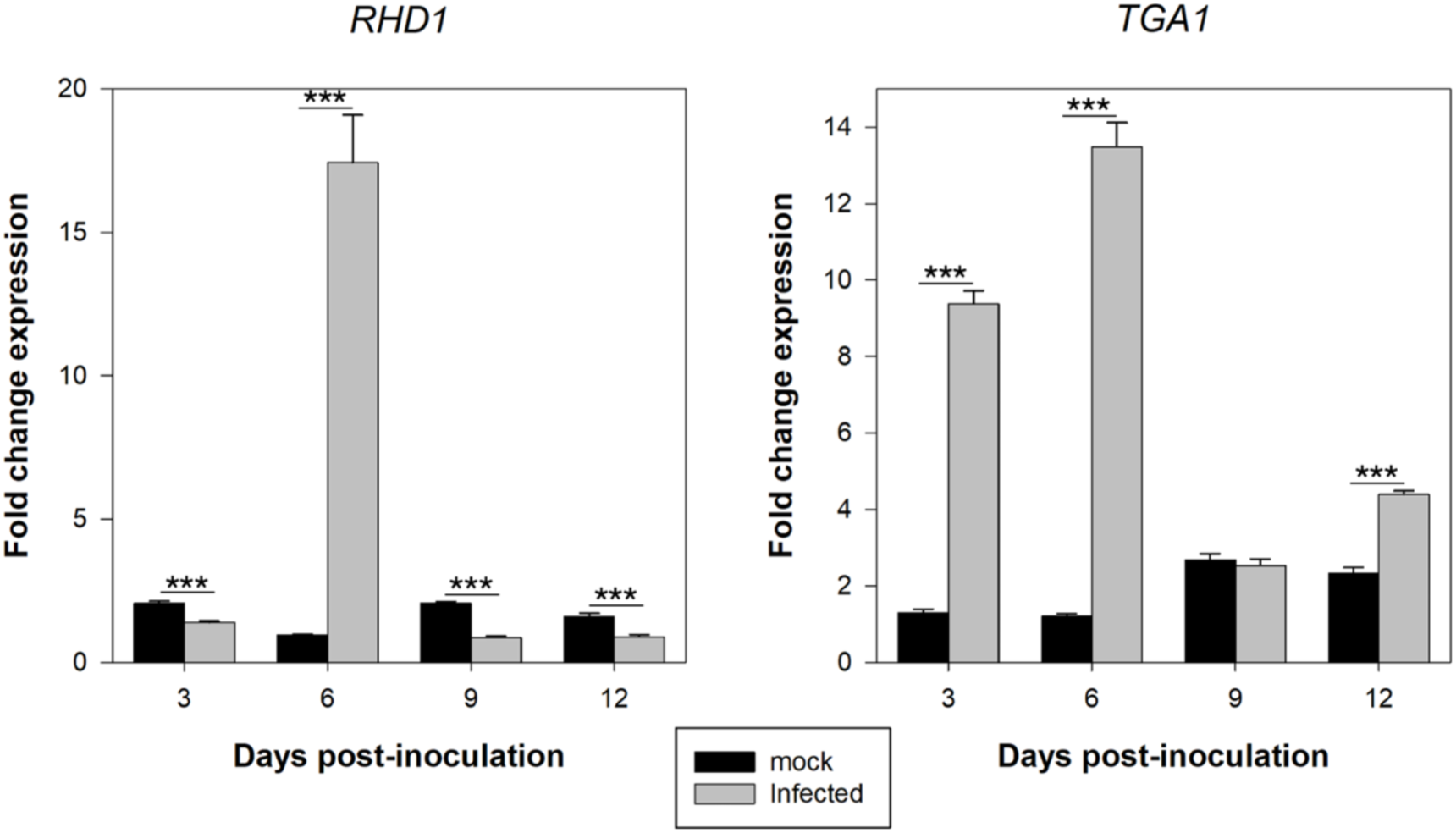
Effect of TuMV VPg^wt^ infection upon expression of VPg direct (*RHD1*) and indirect (*TGA1*) interactors. Significance levels (sequential Bonferroni *post hoc* tests: *** *P* < 0.001.

In the case of *RHD1*, overall effects were observed between infected and non-infected plants (GLM: χ² = 28.568, 1 d.f., *P* < 0.001, 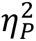 = 0.814) and across the experimental time (χ² = 87.164, 3 d.f., *P* < 0.001, 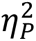 = 0.942). More notably, a highly significant interaction between these two factors was detected (χ² = 102.188, 3 d.f., *P* < 0.001, 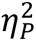 = 0.953). This interaction is explained by gene expression in mock-inoculated plants being consistently low and stable over time. In sharp contrast, expression in infected plants was significantly lower (1.5- to 2.4-fold range; *P* < 0.001) at all time points except at 6 dpi when *RHD1* expression surged 20-fold higher in infected plants (*P* < 0.001). Such pulse-like gene expression, is well suited for coordinating innate immune antiviral responses, fine-tuning cellular processes without committing to prolonged changes in gene activity (*e.g*., Czerkies et al. 2018).

Concerning the effect of infection on *TGA1* expression, both infection status (χ² = 100.553, 1 d.f., *P* < 0.001, 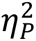 = 0.980) and dpi (χ² = 39.247, 3 d.f., *P* < 0.001, 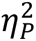) had significant effects on *TGA1* expression levels. These effects are not independent (χ² = 89.968, 3 d.f., *P* < 0.001, 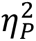 = 0.975), as infected plants exhibit enhanced expression over the time course. Maximum differences with non-infected plants were observed at the two earlier time points (7.2- and 11.2-fold higher, respectively; *P* < 0.001), followed by a reduction to non-significant differences at 9 dpi (*P* = 0.570) or a 1.9-fold higher expression at the latest sampled time point (*P* < 0.001). This biphasic step-down expression profile reflects a dynamic regulation of *TGA1* gene activity. The initial high expression phase would be associated with an acute response to the initial infection, while the reduction may represent a shift to a more sustained level of expression compatible with TuMV overcoming defenses and stablishing a successful infection.

TGA1 regulates plant defense in a manner independent of NPR1 (Shearer et al. 2012). It accomplishes this by inducing the expression of key transcription factors, such as *SYSTEMIC ACQUIRED RESISTANCE DEFICIENT 1* and *CALMODULIN-BINDING PROTEIN 60g*. These transcription factors, in turn, target various essential regulators of plant defense, including enzymes crucial for the synthesis of SA (Sun et al. 2018). Upon a pathogen attack, SA levels surge, triggering two distinct responses: (*i*) the activation of *NPR1*, leading to the expression of pathogenesis-related genes critical for reinforcing systemic acquired resistance (SAR) against subsequent pathogen assaults (Zavaliev and Dong 2023) and (*ii*) the enhancement of RNA-silencing antiviral defense (Alamillo, Saénz and García 2006). Notably, despite the unaffected SAR in *tga1* plants, they exhibit increased susceptibility to *Pseudomonas syringae* (Shearer et al. 2012). However, these plants paradoxically display heightened resistance to TuMV infection (Martínez et al. 2023). This discrepancy raises intriguing questions about whether the activation of *TGA1* during infection is a specific plant response or a consequence of a virus-induced overexpression.

### 3.3. *eif(iso)4e* resistance to TuMV infection is not affected by the three VPg alleles

Sets of twenty plants of genotypes *eif(iso)4e* and wild-type were individually inoculated with the three different VPg alleles. Daily records were kept for both the number of infected plants and the severity of symptoms. A notable initial observation was that none of the 20 *eif(iso)4e* plants exhibited any infection symptoms, irrespective of the VPg allele carried by the inoculated TuMV. The possibility of this negative result being attributed to failed inoculation trials was considered. To eliminate this potential explanation, an additional 68 wild-type plants were inoculated with TuMV-VPg^wt^. With a sample size of 88 wild-type plants, the probability of infecting a fully susceptible wild-type plant with TuMV-VPg^wt^ was estimated based on the number of positive events (86) observed after inoculating all the wild-type plants. Using the LaPlace point estimator and the adjusted Wald 95% confidence interval method, the success rate was calculated to be 0.967 (0.916, 0.999). Therefore, even in the less favorable situation (*i.e*., the lower limit of the confidence interval), the likelihood of failing 20 independent inoculation events of fully susceptible plants with a wild-type virus would be given by the Bernoulli process (1 – 0.916)^20^ = 3.06·10^−22^.

The same rationale was applied to the two mutant VPg alleles, where 20 out of 20 inoculated wild-type plants resulted in symptomatic infections. In this case, the LaPlace estimation of success rate was 0.955 (0.859, 1.000), and the Bernoulli measure for the less favorable situation was 1.007·10^−17^. Consequently, we can confidently conclude that *eif(iso)4e* plants are fully resistant to TuMV YC5 infection, regardless of the VPg allele carried by the virus.

Many recessive resistance genes against potyviruses described in wild and crop species encode members of the eIF(iso)4E and eIF(iso)4G families (Truninger and Aranda 2009). Here, we have observed that TuMV YC5 replication strictly depends on the interaction between VPg and a functional eIF(iso)4E. The complete resistance observed for *eif(iso)4e* plants suggests its function cannot be complemented by other translation initiation factors. By contrast, Navarro et al. (2022) were able of experimentally evolving lineages derived from the YC5 isolate in *A. thaliana* mutants *i4g1* (deficient for eIF(iso)4G1), observing different amino acid substitutions in VPg associated with significant fitness increases. These results confirm that eIF(iso)4G is not as essential as eIF(iso)4E for TuMV infection or that its function can be complemented by other translation initiation factors.

### 3.4. Disease progression in *rhd1* mutant plants depends on the VPg allele carried by TuMV

In stark contrast to *eif(iso)4e* plants, *rhd1* plants exhibited variable responses to infection with the three TuMV variants. Firstly, concerning changes in infection rate (Fig. 3A), the mean time to produce visible symptoms was the same for the three viral variants in wild-type plants (log-rank test: χ² = 5.806, 2 d.f., *P* = 0.055), averaging 6.817 ±0.151 dpi. However, significant differences among VPg alleles were observed in *rhd1* plants (χ² = 5.806, 2 d.f., *P* = 0.002): TuMV-VPg^wt^ produced visible symptoms at 7.000 ±0.218 dpi (±1 SE), significantly later than TuMV-VPg^D113G^ (6.700 ±0.242; χ² = 3.881, 1 d.f., *P* = 0.049) and TuMV-VPg^R118H^ (5.950 ±0.170; χ² = 16.028, 1 d.f., *P* < 0.001), although the difference between both mutants was not statistically significant (χ² = 3.839, 1 d.f., *P* = 0.050). Therefore, we conclude that VPg^D113G^ and VPg^R118H^ accelerate the appearance of disease symptoms in plants deficient for RHD1.

**Figure 3.**
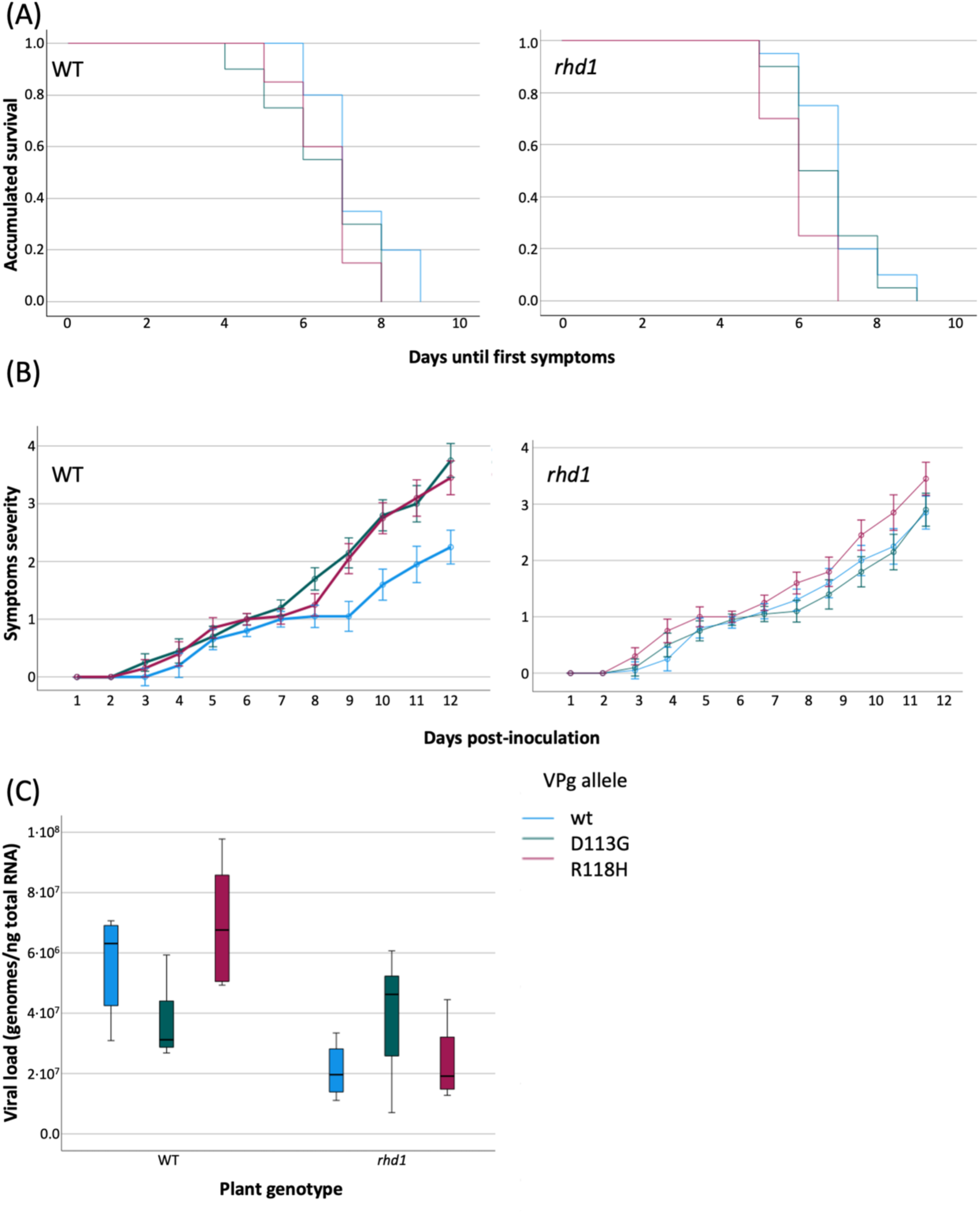
(A) Survival curves representing the number of non-symptomatic plants along time. (B) Progression of disease severity in wild-type and *rhd1* plants infected with TuMV expressing the three different VPg alleles. Error bars represent 95% CI. (C) Differences in viral load. Boxes represent interquartile ranks, horizontal lines the median and error bars represent 95% CI. WT: wild-type plants.

Secondly, Fig. 3B illustrates the disease progression curves for the three virus variants in both wild-type and *rhd1* plants. In wild-type plants, large and significant inter-individual differences were observed between the three VPg alleles (*F*_2,57_ = 45.487, *P* < 0.001, 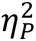 = 0.615), as well as substantial intra-individual differences between dpi (*F*_3.933,224.203_ = 502.504, *P* < 0.001, 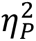 = 0.898) and the interaction between intra- and inter-individual factors (*F*_7.867,224.203_ = 13.586, *P* < 0.001, 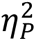 = 0.323). Overall, TuMV-VPg^wt^ induced milder symptoms (0.879 ±0.043) compared to viruses expressing the two mutant VPg variants (1.417 ±0.043 for VPg^D113G^ and 1.338 ±0.043 for VPg^R118H^; pairwise comparisons: *P* < 0.001). The progression of symptoms was similar for the three variants during the first 7 dpi but diverged between VPg^wt^ and the two mutants thereafter. The two mutant VPg variants were indistinguishable throughout the entire time course of the experiment (*P* = 0.595). In the case of *rhd1* plants, inter-individual effects were large and significant (*F*_2,57_ = 8.322, *P* < 0.001, 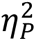 = 0.226), as were intra-individual effects of dpi (*F*_2.790,159.046_ = 293.746, *P* < 0.001, 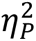 = 0.837), but not the interaction of the two factors (*F*_5.581,159.046_ = 2.090, *P* = 0.062, 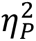 = 0.068). This suggests that differences among virus variants remained constant over time. Overall, indistinguishable symptoms were induced by TuMV-VPg^wt^ and TuMV-VPg^D113G^ (1.092 ±0.059 and 1.058 ±0.059, respectively; *P* = 1.000), but both were significantly milder than those induced by TuMV-VPg^R118H^ (1.371 ±0.059; *P* ≤ 0.005).

Thirdly, we assessed the efficiency of TuMV carrying different VPg alleles in terms of viral accumulation at 12 dpi. Fig. 3C illustrates the comparison between the two plant genotypes and the three VPg alleles. The two orthogonal main factors had a significant effect on viral load, although the magnitude of the effect was small among VPg alleles (plant genotype: χ² = 226.733, 1 d.f., *P* < 0.001, 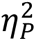 = 0.459; VPg allele: χ² = 75.113, 2 d.f., *P* < 0.001, 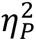 = 0.065). Nonetheless, the interaction between both factors was highly significant and of large magnitude (χ² = 196.689, 2 d.f., *P* < 0.001, 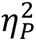 = 0.353), suggesting that the effect on viral load of the VPg alleles was strongly dependent on the expression of *RHD1*. As illustrated in Fig. 3C, viral load was reduced, on average, by 47.93% in *rhd1* plants (*P* ≤ 0.018 in the three pairwise comparisons). Focusing on wild-type plants, TuMV-VPg^R118H^ showed the highest log-viral load (7.839 ±6.200; ±1 SE), followed by TuMV-VPg^wt^ (7.754 ±6.122) and TuMV-VPg^D113G^ (7.563 ±5.992). However, in *rhd1* plants, the situation reversed, and the highest log-viral load was observed for TuMV-VPg^D113G^ (7.597 ±6.515), followed by TuMV-VPg^wt^ (7.329 ±5.824) and TuMV-VPg^R118H^ (7.375 ±5.997), showing very similar accumulations (*P* ≤ 0.031 in the three pairwise comparisons).

At first glance, the observation of earlier and more severe symptoms in *rhd1* plants suggests that the RHD1 protein plays an antiviral role in *A. thaliana*. However, higher TuMV accumulation was noted in wild-type plants. To reconcile these two observations, one could argue that symptoms may not necessarily correlate with viral accumulation if *RHD1* enhances the plant’s tolerance to viral infection (Pagán and García-Arenal 2020). In such a scenario, wild-type plants expressing *RHD1* might exhibit higher viral accumulation while maintaining mild symptoms. Conversely, if *RHD1* expression is knocked out, even low viral accumulation could rapidly result in more severe symptoms. Interestingly, the level of tolerance appears to be influenced by VPg alleles: viruses carrying the wild-type and R118H alleles demonstrate larger reductions in virus accumulation, whereas viruses carrying the generalist D113G allele are less affected (Fig. 3C).

Together with RMB1 and ROS1-ASSOCIATED WD40 DOMAIN-CONTAINING PROTEIN (RWD40), RHD1 is part of the RWD40 complex, which regulates active DNA demethylation by specifically directing ROS1 DNA demethylase to specific target sequences in the *A. thaliana* genome (Liu et al. 2021). The absence of expression of any of the components of the RWD40 complex results in ROS1 mislocalization, hypermethylation of targeted genomic DNA regions and hypersusceptibility to the hemibiotrophic bacterial pathogen *Pseudomonas syringae*, suggesting a role of the RWD40 DNA demethylation complex in pathogen defense (Liu et al. 2021). One of these regions is localized in the *ROS1* promoter, leading to slightly increased *ROS1* expression. Therefore, in *rhd1* plants, *ROS1* expression is upregulated but its methylated target sequences are not recognized and become hypermethylated in compared to wild-type plants. In addition, TuMV infection also downregulates the expression of *ROS1* (Corrêa et al. 2020), in agreement with the results shown for wild-type plants in Fig. 2 for most dpi. The sharp *RHD1* induction observed during TuMV infection might be involved in temporally downregulating *ROS1* expression and the subsequent temporal increase in the methylation rate of specific genome sequences, thereby altering gene expression. Moreover, the pulse-like expression of *RHD1* and the drop in expression in *TGA1* observed after 6 dpi (Fig. 2) coincides with the slow-down in symptoms progression observed for TuMV-VPg^wt^, which stay subsequently milder (roughly speaking in the range 1 to 2). The two VPg mutants escape from this control, resulting in worsened symptoms.

### 3.5. Integrating interaction results with structural analyses

To better understand these differences in interactions, we modeled the interactions of VPg with eIF(iso)4E (Fig. 4) and RHD1 (Fig. 5). First, we modeled the TuMV-VPg/eIF(iso)4E complex using a docking procedure guided by experimental data (Supplementary Fig. S2). Several lines of evidence suggest that the proposed model is close to the real structure: (*i*) the contact surface involves the central domain of VPg and the cap-binding region of eIF(iso)4E, in agreement with experimental data; (*ii*) conserved residue R103 in the central domain of VPg binds to eIF(iso)4E in a position equivalent to the guanine base in the m^7^G cap, (*iii*) acidic residues in the segment comprising the α1-α2 loop bind to a region close to the phosphate binding region in eIF(iso)4E, (*iv*) experimental data on residues involved int eIF4E - VPg interactions are mostly clustered in the docking interaction surface (Grzela et al. 2006; Roudent-Tavert et al. 2007; Ashby et al. 2012; Svanella-Dumas et al. 2014; Coutinho de Oliveira et al. 2019). Our model also provides a plausible explanation of the effect of the R118H and D113G substitutions on the eIF(iso)4E - VPg interaction (Fig 4). R118 and D113 are located in a small helical segment predicted to bind to a variable region of complementary charge in eIF(iso)4E in the m^7^G cap binding pocket (Fig. 4). Both mutants are predicted to have a significant effect on the net charge with respect to the VPg^wt^ (pI ≈ 7.23), with D113G making the protein more positive (pI ≈ 8.02) and R118H more negative (pI ≈ 6.74). Therefore, we postulate that these pI changes in the R118H and D113G mutants remove favorable ionic interactions, resulting in the observed reduction in the stability of the eIF(iso)4E - VPg complex.

**Figure 5.**
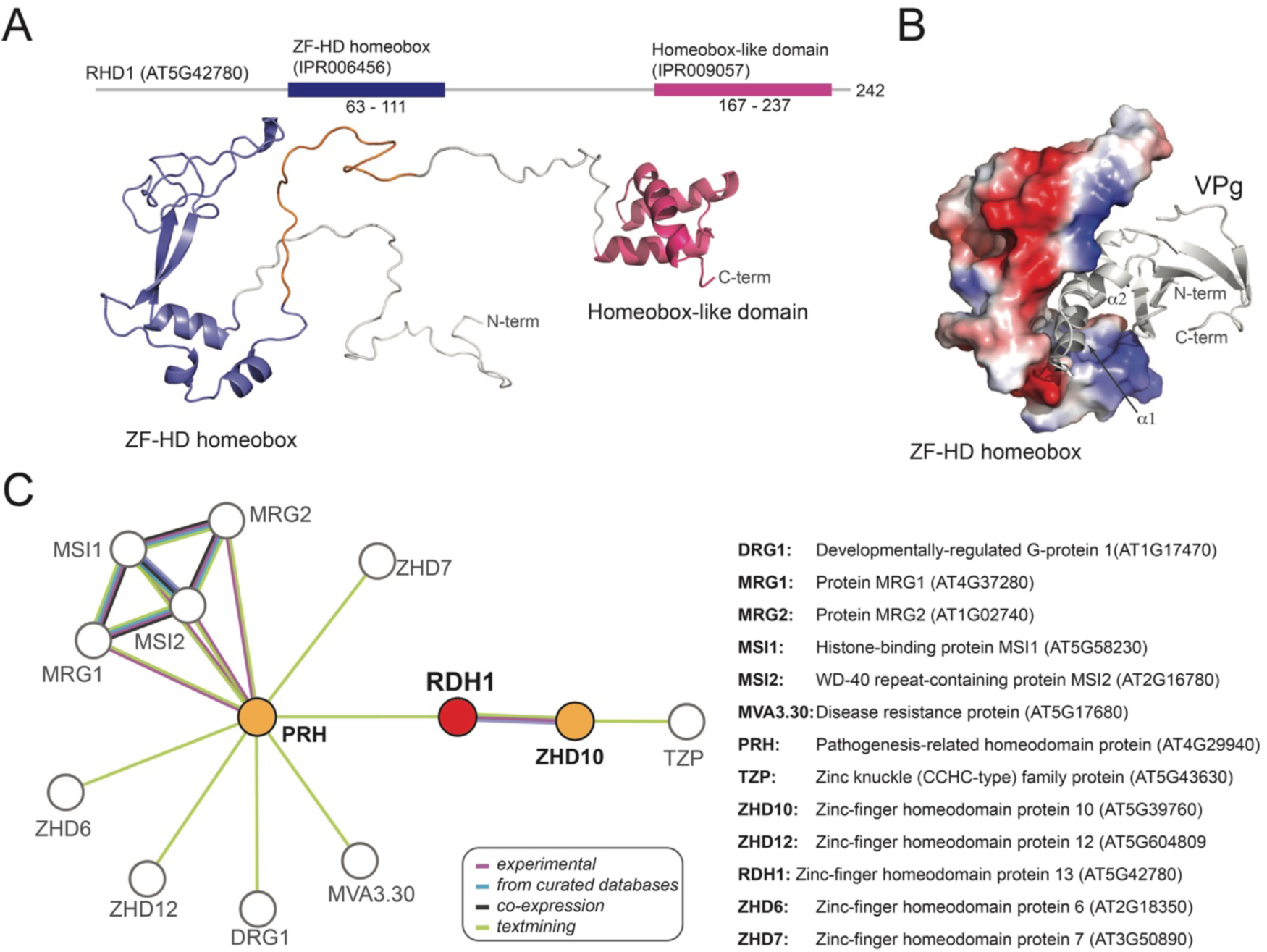
Structural properties of the RHD1 protein from *A. thaliana*. (A) The AlphaFold modeling and sequences analysis reveal that this protein contains Zn-finger (ZF-HD) and a C-terminal homeobox domain, and unstructured N-terminal and central regions. (B) Global docking of VPg against the homeobox domains of RHD1using ROSETTA revealed a potential binding site comprising the central domain of VPg and a crevice in the ZF-HD homeobox region. (C) PPI network of the RHD1 with other *A. thaliana* proteins according to STRING database.

Our model also explains the resistance of *eif(iso)4e* plants against the three TuMV mutants. Potyviruses selectively require different eIF4E paralogs to establish infection (Duprat et al. 2002; Sato et al. 2005; Nicaise et al. 2007; Gómez et al. 2019). In *A. thaliana*, lettuce mosaic virus (LMV), TEV and TuMV use eIF(iso)4E, while other potyviruses, such as clover yellow vein virus and PVY, use eIF4E (Sato et al. 2005; Zafirov et al. 2023). Structurally, one of the most salient differences between eIF(iso)4E and its paralogs eIF4E and nCBP is the presence of a bulkier segment in the VPg binding region. We propose that due to this steric effect, neither eIF4E nor nCBP can substitute for eIF(iso)4E in the *eif(iso)4e* plants (Fig. 4).

We also used docking to identify a potential VPg binding pocket in the RHD1 protein from *A. thaliana* (Fig. 5). A prediction of structural domains revealed that this protein contains an N-proximal Zn finger domain, a C-terminal homeobox domain and two unstructured segments comprising the N-terminal and central regions (Fig. 5A). Assuming that binding occurs in the homeobox regions, we performed a global docking of VPg against the homeobox domains using a refined AlphaFold model of RHD1 (Fig. 5B). The lowest energy complex fitted the VPg core domain within a crevice separating the helical and beta-stranded regions of the ZF-HD homeobox. This region comprises domains of positive and negative charge that might be affected by the ionization state of the VPg protein.

Fig. 5C shows the RHD1 PPIs network generated with STRING. Interestingly, RHD1 interactors mostly belong to the ZF-HD-containing family protein of transcription factors. The direct interactor PATHOGENESIS RELATED HOMEODOMAIN PROTEIN A (PRHA) is a hub that bridges ZF-HDs involved in the regulation of different aspects of plant development and of circadian clock, with MVA3.30, a disease resistance protein of the TIR-NBS-LRR class family, and with MORPH RELATED GENE 1 (MRG1) and 2 (MRG2), both readers of H3K4m3/H3K36m3 methylation marks. Taken together, these interactions suggest that VPg affects epigenetic regulations by directly interacting with RHD1 and indirectly affecting its interaction with other transcription factors and with components of the histone methylation machinery. The role of such epigenetic pathways in TuMV infection has been already stablished (Corrêa et al. 2020; Navarro et al. 2022) and might contribute to the enhanced tolerance to infection of plants accumulating RHD1.

## 4. Concluding remarks

The results shown above shed light on the complex interplay between TuMV VPg variants, host proteins, and plant defense response, providing valuable insights into the molecular mechanisms underlying virus-host interactions and disease progression in *A. thaliana*. The three different VPg alleles used in our study show differential interactions with plant proteins. Infected plants show altered expression of key genes *eIF(iso)4E*, *RHD1* and *TGA1*. Mutant plants *eif(iso)4e* show complete resistance to TuMV infection regardless of the VPg allele, confirming its well described essential role in virus replication. Finally, different VPg alleles affect disease progression in *rhd1* mutant plants. VPg^D113G^ and VPg^R118H^ accelerate symptom appearance, while VPg^wt^ induces milder symptoms. This observation suggests that RHD1 modulates the tolerance to viral infection, likely via epigenetic modifications.

## Data availability

All raw data are available at https://git.csic.es/SFElena/tumv-evolution-in-a.-thaliana-genotypes-with-deficient-immune-responses.

## Supplementary data

Supplementary data is available at *Virus Evolution* online.

## Acknowledgments

We thank Francisca de la Iglesia and Paula Agudo for excellent technical assistance and the rest of the EvolSysVir lab members for fruitful discussions.

---

**Supplementary Figure S1.**
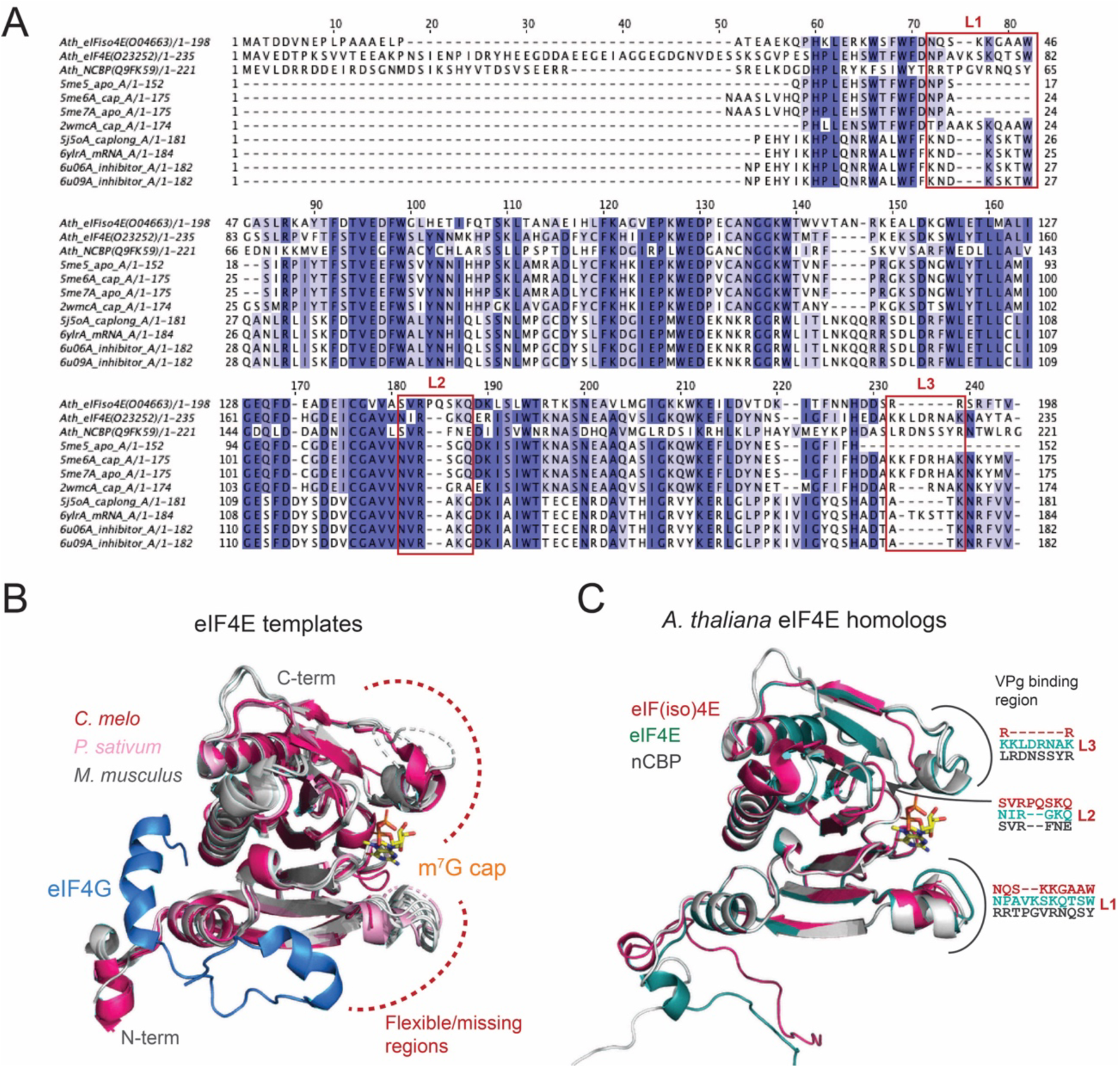
Sequence and structural analysis of the eIF(iso)4E structure from *A. thaliana*. (A) Sequence alignment of *A. thaliana* eIF(iso)4E homologs and the eIF4E sequences used as structural templates. Variable loop regions predicted to be involved in the interaction with VPg are enclosed in red boxes. (B) Superposition of the structural templates used in this work showing position of the m^7^G and eIF4G binding regions. (C) Superposition of the eIF(iso)4E, eIF4E and nCBP model from *A. thaliana* highlighting sequence and structural differences in the VPg-binding loops.

**Supplementary Figure S2.**
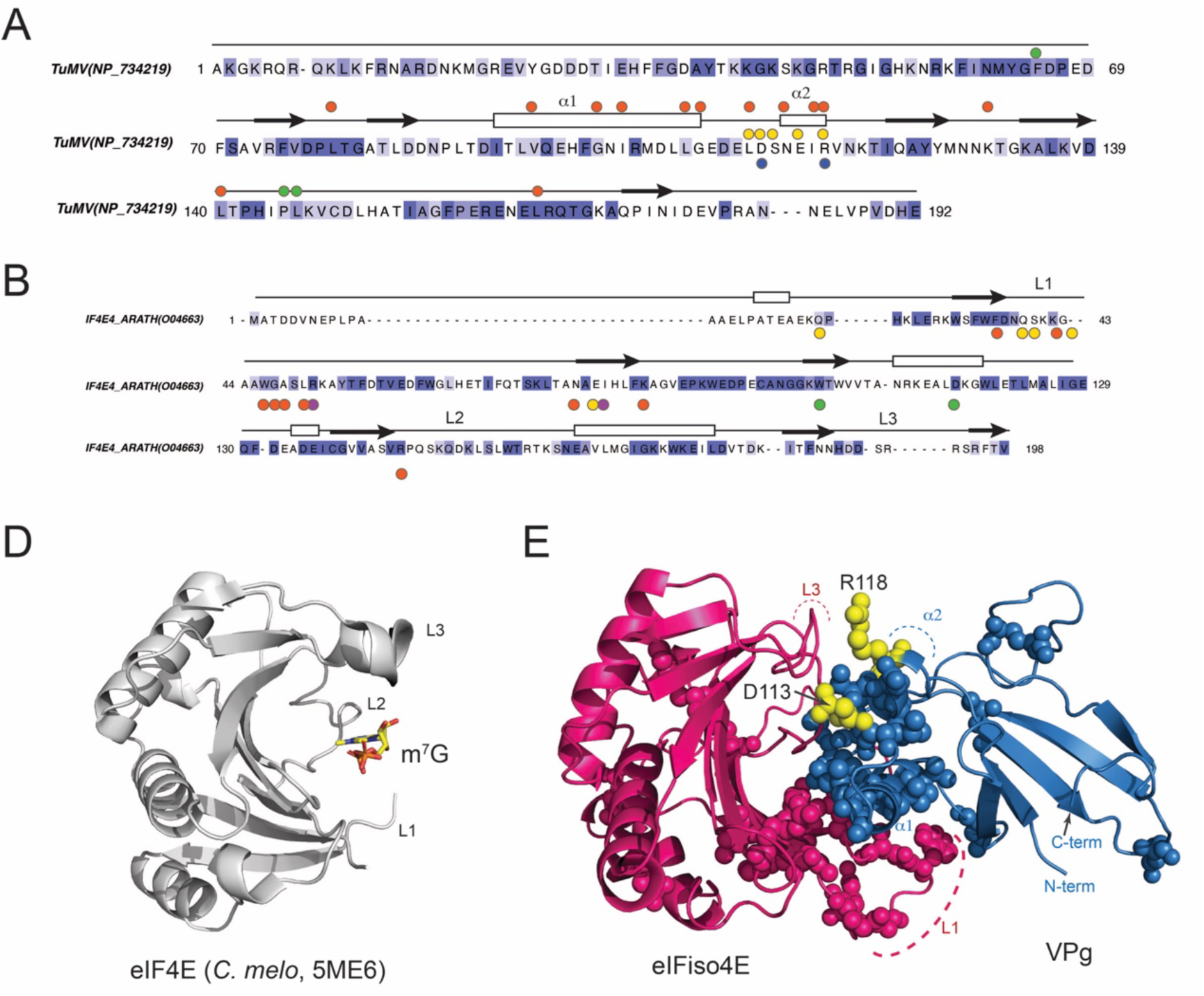
Constraints used in the construction of the eIF(iso)4E - TuMV VPg model. (A) Identification of potential eIF(iso)4E interacting residues in the VPg based on a multiple sequence alignment of VPg protein from LMV, PVY, TEV, and TuMV. Filled circles indicate regions shown to be involved in the interaction with eIF4E/eIF(iso)4E: Coutinho de Olivera et al. (2019) (orange), Roudet-Tavert et al. (2007) (purple), Pérez et al. (2012) (yellow), Svanella-Dumas (2014) (green), and this study (blue). (B) dentification of potential VPg interacting residues in the eIF(iso)4E from *A. thaliana*. In this case, filled circles correspond to interaction data from Coutinho de Olivera *et al*. (2019) (orange), German-Retana et al. (2008) (purple), Pérez et al. (2012) (yellow), Ashby et al. (2011) (green), and Monzingo et al. (2007) (blue). (C) Structure of the eIF4E from *C. melo* in complex with m^7^G, illustrating the position of the putative VPg-binging loops. (D) Cartoon representation of the predicted eIF(iso)4E - VPg complex derived from the interaction data presented in panels A and B. Interacting residues are shown as spheres. Residues D113 and R118 are shown in yellow.

**Supplementary Figure S3.**
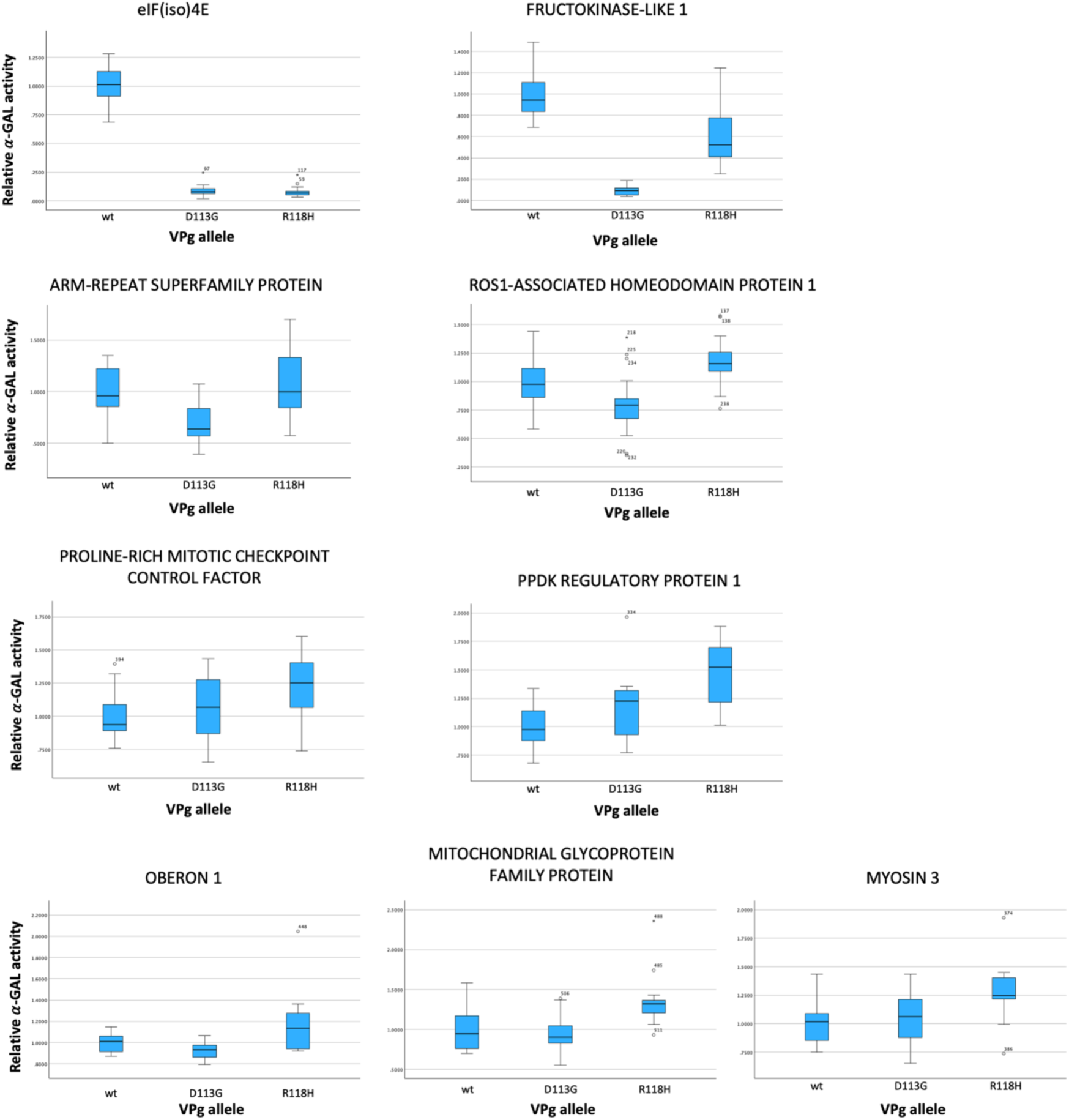
Distribution of affinity binding values estimated for the nine host interactors for which a significant difference between the mutant VPg alleles and the VPg^wt^ one has been observed.

## References

Agudelo-Romero, P. et al. (2008) ‘Virus adaptation by manipulation of host’s gene expression’, PLoS ONE, 3: e2397.

Aguirre, J., and Guantes R. (2023) ‘Virus-host protein co-expression networks reveal temporal organization and strategies of viral infection’, iScience, 26: 108475.

Ahmed, H. et al. (2018) ‘Network biology discovers pathogen contact points in host protein-protein interactomes’, Nature Communications, 9: 2312.

Alamillo, J.M., Saénz, P., and García, J.A. (2006) ‘Salicylic acid-mediated and RNA- silencing defense mechanisms cooperate in the restriction of systemic spread of plum pox virus in tobacco’, Plant Journal, 48: 217–27.

Amari, K. et al. (2014) ‘Myosins VIII and XI play distinct roles in reproduction and transport of *Tobacco mosaic virus*’, PLoS Pathogens, 10: e1004448.

Ambrós, S. et al. (2024) ‘Phenotypic and genomic changes during turnip mosaic virus adaptation to *Arabidopsis thaliana* mutants lacking epigenetic regulatory factors’, Evolution, 28: 69–85.

Ashby, J. A. et al. (2011) ‘Structure-based mutational analysis of eIF4E in relation to *sbm1* resistance to pea seed-borne mosaic virus in pea’, PLoS ONE, 6: e15873.

Ayme, V. et al. (2006) ‘Different mutations in the genome-linked protein VPg of potato virus Y confer virulence on the *pvr2^3^* resistance in pepper’, Molecular Plant-Microbe Interactions, 19: 557–63.

Belshaw, R., Pybus, O. G., and Rambaut, A. (2007) ‘The evolution of genome compression and genomic novelty in RNA viruses’, Genome Research, 17: 1496–504.

Bosque, G. et al. (2014) ‘Topology analysis and visualization of *Potyvirus* protein-protein interaction network’, BMC Systems Biology, 8: 129.

Boyes, D. C. et al. (2001) ‘Growth stage-based phenotypic analysis of Arabidopsis: a model for high throughput functional genomics in plants’, Plant Cell, 13: 1499–510.

Buschmann, H. et al. (2004) ‘Helical growth of the *Arabidopsis* mutant *tortifolia1* reveals a plant-specific microtubule-associated protein’, Current Biology, 14: 1515–21.

Butković, A. et al. (2021) ‘A genome-wide association study identifies *Arabidopsis thaliana* genes that contribute to differences in the outcome of infection with two turnip mosaic potyvirus strains that differ in their evolutionary history and degree of host specialization’, Virus Evolution, 7: veab063.

Cervera, H. et al. (2018) ‘Viral fitness correlates with the magnitude and direction of the perturbation induced in the host’s transcriptome: the tobacco etch potyvirus-tobacco case study’, Molecular Biology and Evolution, 35: 1599–615.

Chastain, C. J. et al. (2007) ‘The pyruvate, orthophosphate dikinase regulatory proteins of Arabidopsis possess a novel, unprecedented Ser/Thr protein kinase primary structure’, Plant Journal, 53: 854–63.

Chaudhury, S. et al. (2011) ‘Benchmarking and analysis of protein docking performance in Rosetta v3.2’. PLoS ONE, 6: e22477.

Chen, C. C. et al. (2003) ‘Identification of turnip mosaic virus isolates causing yellow stripe and spot on calla lily’, Plant Disease, 87: 901–5.

Conway, P. et al. (2014) ‘Relaxation of backbone bond geometry improves protein energy landscape modeling’, Protein Science, 23: 47–55.

Corrêa, R. L. et al. (2020) ‘Viral fitness determines the magnitude of transcriptomic and epigenomic reprogramming of defense responses in plants’, Molecular Biology and Evolution, 37: 1866–81.

Coutinho de Oliveira, L., et al. (2019) ‘Structural studies of the eIF4E-VPg complex reveal a direct competition for capped RNA: implications for translation’, Proceedings of the National Academy of Sciences of the USA, 116: 24056–65.

Czechowski, T. et al. (2005) ‘Genome-wide identification and testing of superior reference genes for transcript normalization in Arabidopsis’, Plant Physiology, 139: 5–17.

Czerkies, M. et al. (2018) ‘Cell fate in antiviral response arises in the crosstalk of IRF, NF-κB and JAK/STAT pathways’, Nature Communications, 9: 493.

De Chassey, B. et al. (2008) ‘Hepatitis C virus infection protein network’, Molecular Systems Biology, 4: 230.

Dominguez, C., Boelens, R., and Bonvin, A. M. (2003) ‘HADDOCK: a protein-protein docking approach based on biochemical or biophysical information’, Journal of the American Chemical Society, 125: 1731–7.

Du, P. et al. (2023) ‘WRKY transcription factors and OBERON histone-binding proteins form complexes to balance plant growth and stress tolerance’, EMBO Journal, 42: e113639.

Duprat, A. et al. (2002) ‘The Arabidopsis eukaryotic initiation factor (iso)4e is dispensable for plant growth but required for susceptibility to potyviruses’, Plant Journal, 32: 927–34.

Dyer, M. D., Murali, T. M., and Sobral, B. W. (2008) ‘The landscape of human proteins interacting with viruses and other pathogens’, PLoS Pathogens, 4: e32.

Edgar, R. C. (2004) ‘MUSCLE: multiple sequence alignment with high accuracy and high throughput’, Nucleic Acids Research, 32: 1792–7.

Eskelin, K. et al. (2011) ‘Potyviral VPg enhances viral RNA translation and inhibits reporter mRNA translation *in planta*’, Journal of Virology, 85: 9210–21.

Gallois, J. L. et al. (2010) ‘Single amino acid changes in the turnip mosaic virus viral genome-linked protein (VPg) confer virulence towards *Arabidopsis thaliana* mutants knocked out for eukaryotic initiation factors eIF(iso)4E and eIF(iso)4G’, Journal of General Virology, 91: 288–93.

German-Retana, S. et al. (2008) ‘Mutational analysis of plant cap-binding protein eIF4E reveals key amino acids involved in biochemical functions and potyvirus infection’, Journal of Virology, 82: 7601–12.

Gilkerson, J. et al. (2012) ‘The plastid-localized pfkB-type carbohydrate kinases FRUCTOKINASE-LIKE 1 and 2 are essential for growth and development of *Arabidopsis thaliana*’, BMC Plant Biology, 12: 102.

Gómez, M. A. et al. (2019) ‘Simultaneous CRISPR/Cas9-mediated editing of cassava eIR4E isoforms nCBP-1 and nCBP-2 reduce cassava brown streak disease symptom severity and incidence’, Plant Biotechnology Journal, 17: 421–34.

González, R., Butković, A., and Elena, S. F. (2019) ‘Role of host genetic diversity for susceptibility-to-infection in the evolution of virulence of a plant virus’, Virus Evolution, 5: vez024.

González, R. et al. (2021) ‘Plant virus evolution under strong drought conditions results in a transition from parasitism to mutualism’, Proceedings of the National Academy of Sciences of the USA, 118: e2020990118.

Grzela, R. et al. (2006) ‘Potyvirus terminal protein VPg, effector of host eukaryotic initiation factor eIF4E’, Biochimie, 88: 887–96.

Hafrén, A., Löhmus, A., and Mäkinen, K. (2015) ‘Formation of potato virus A-induced RNA granules and viral translation are interrelated processes required for optimal virus accumulation’, PLoS Pathogens, 11: e1005314.

Hillung, J. et al. (2014) ‘Experimental evolution of an emerging plant virus in host genotypes that differ in their susceptibility to infection’, Evolution, 68: 2467–80.

Honorato, R. V. et al. (2021) ‘Structural biology in the clouds: the WeNMR-EOSC ecosystem’, Frontiers in Molecular Biosciences, 8: 729513.

Jumper, J. et al. (2021) ‘Highly accurate protein structure prediction with AlphaFold’, Nature, 596: 583–9.

Liu, P. et al. (2021) ‘A novel protein complex that regulates active DNA demethylation in Arabidopsis’, Journal of Integrative Plant Physiology, 63, 772–86.

Mahmoudabadi, G, and Phillips, R. (2018) ‘A comprehensive and quantitative exploration of thousands of viral genomes’, eLife, 7: e31955.

Mandell, D. J. et al. (2009) ‘Sub-angstrom accuracy in protein loop reconstruction by robotics-inspired conformational sampling’, Nature Methods, 6: 551–2.

Martínez, F. et al. (2016) ‘Interaction network of tobacco etch potyvirus NIa protein with the host proteome during infection’, BMC Genomics, 17: 87.

Martínez, F. et al. (2023) ‘A binary interaction map between turnip mosaic virus and *Arabidoposis thaliana* proteomes’, Communications Biology, 6: 28.

Melero, I., González, R. and Elena, S.F. (2023) ‘Host developmental stages shape the evolution of plant RNA virus’, Philosophical Transactions of the Royal Society B: Biological Sciences, 378: 20220005.

Menche, J. et al. (2015) ‘Uncovering disease-disease relationships through the incomplete interactome’, Science, 347: 841.

Miras, M. et al. (2017) ‘Structure of eIF4E in complex with an eIF4G peptide supports a universal bipartite binding mode for protein translation’, Plant Physiology, 174: 1476–91.

Monzingo, A. F. et al. (2007) ‘The structure of eukaryotic translation initiation factor-4E from wheat reveals a novel disulfide bond’, Plant Physiology, 143: 1504–18.

Moury, B. et al. (2004) ‘Mutations in potato virus Y genome-linked protein determine virulence towards recessive resistances in *Capsicum annuum* and *Lycopersicon hirsutum*’, Molecular Plant-Microbe Interactions, 17: 322–9.

Mukhtar, M. S. et al. (2011) ‘Independently evolved virulence effectors converge onto hubs in a plant immune system network’, Science, 333: 596–601.

Navarro, R. et al. (2022) ‘Defects in plant immunity modulate the rates and patterns of RNA virus evolution’, Virus Evolution, 8: veac059.

Nicaise, V. et al. (2007) ‘Coordinated and selective recruitment of eIF4E and eIF4G factors for potyvirus infection in *Arabidopsis thaliana*’, FEBS Letters, 581: 1041–6.

Pagán, I., and García-Arenal, F. (2020) ‘Tolerance of plants to pathogens: an unifying view’, Annual Review of Phytopathology, 58: 77–96.

Peremyslov, V. V. et al. (2008) ‘Two class XI myosins function in organelle trafficking and root hair development in Arabidopsis’, Plant Physiology, 146: 1109–16.

Pérez, K. et al. (2012) ‘Tobacco etch virus infectivity in *Capsicum* spp. is determined by a maximum of three amino acids in the viral virulence determinant VPg’, Molecular Plant Microbe Interactions, 25: 1562–73.

Phillips, J. C. et al. (2020). ‘Scalable molecular dynamics on CPU and GPU architectures with NAMD’, Journal of Chemical Physics, 153: 044130.

Pichlmair, A. et al. (2012) ‘Viral immune modulators perturb the human molecular network by common and unique strategies’, Nature, 487: 486–90.

Roudet-Tavert, G. et al. (2007) ‘Central domain of a potyvirus VPg is involved in the interaction with the host translation initiation factor eIF4E and the viral protein HCPro’, Journal of General Virology, 88: 1029–33.

Rydzik, A. M. et al. (2017) ‘mRNA cap analogues substituted in the tetraphosphate chain with CX2: identification of O-to-CCl2 as the first bridging modification that confers resistance to decapping without impairing translation’. Nucleic Acids Research, 45: 8661–75.

Sato, M. et al. (2005) ‘Selective involvement of members of the eukaryotic initiation factor 4E family in the infection of *Arabidopsis thaliana* by potyviruses’, FEBS Letters, 579: 1167–71.

Shearer, H.L. et al. (2012) ‘Arabidopsis clade I TGA transcription factors regulate plant defenses in an NPR1-independent fashion’, Molecular Plant Microbe Interactions, 25: 1459–68.

Spivak, M. et al. (2023) ‘VMD as a platform for interactive small molecule preparation and visualization in quantum and classical simulations’, Journal of Chemical Information and Modeling, 63: 4664–78.

Sun, T. et al. (2018) ‘TGACG-BINDING FACTOR 1 (TGA1) and TGA4 regulate salicylic acid and pipecolic acid biosynthesis by modulating the expression of *SYSTEMIC ACQUIRED RESISTANCE DEFICIENT 1* (*SARD1*) and *CALMODULIN-BINDING PROTEIN 60g* (*CBP50g*)’, New Phytologist, 217: 344–54.

Svanella-Dumas, L. et al. (2014) ‘Adaptation of lettuce mosaic virus to *Catharanthus roseus* involves mutations in the central domain of the VPg’, Molecular Plant Microbe Interactions, 27: 491–7.

Szklarczyk, D. et al. (2023) ‘The STRING database in 2023: protein-protein association networks and functional enrichment analyses for any sequenced genome of interest’, Nucleic Acids Research, 50: gkac1000.

Tarazona, A., Forment, J., and Elena, S. F. (2020) ‘Identifying early warning signals for the sudden transition from mild to severe tobacco etch disease by dynamical network biomarkers’, Viruses, 12: 16.

Tisoncik-Go, J. et al. (2016) ‘Integrated omics analysis of pathogenic host responses during pandemic H1N1 influenza virus infection: the crucial role of lipid metabolism’, Cell Host Microbe, 19: 254–66.

Truninger, V., and Aranda M.A. (2009) ‘Recessive resistance to plant viruses’, Advances in Virus Research, 75: 119–59.

Uetz, P. et al. (2006) ‘Herpesviral protein networks and their interaction with the human proteome’, Science, 311: 239–42.

Van Zundert, G. C. P. et al. (2016) ‘The HADDOCK2.2 web server: user-friendly integrative modeling of biomolecular complexes’, Journal of Molecular Biology, 428: 720–5.

Wan, X. et al. (2020) ‘Discovery of lysine-targeted eIF4E inhibitors through covalent docking’. Journal of the American Chemical Society, 142: 4960–4.

Wang, C., Bradley, P., and Baker, D. (2007). ‘Protein-protein docking with backbone flexibility’, Journal of Molecular Biology, 373: 503–19.

Webb, B., and Sali, A. (2016) ‘Comparative protein structure modeling using MODELLER’. Current Protocols in Bioinformatics, 54: 5.6.1-37.

Wojcik, R. et al. (2021) ‘Novel N7-arylmethyl substituted dinucleotide mrna 5’ cap analogs: synthesis and evaluation as modulators of translation’, Pharmaceutics, 13: 1941.

Zafirov, D. et al. (2023) ‘Arabidopsis *eIF4E1* protects the translational machinery during TuMV infection and restricts virus accumulation’, PLoS Pathogens, 19: e1011417.

Zavaliev, R., and Dong, X. (2023) ‘NPR1, a key immune regulator of plant survival under biotic and abiotic stresses’, Molecular Cell, 84: 131–41.

